# Identification and Characterization of Human Monoclonal Antibodies for Immunoprophylaxis Against Enterotoxigenic Escherichia coli Infection

**DOI:** 10.1101/320044

**Authors:** Serena Giuntini, Matteo Stoppato, Maja Sedic, Monir Ejemel, Jessica R. Pondish, Danielle Wisheart, Zachary A. Schiller, William D. Thomas, Eileen M. Barry, Lisa A. Cavacini, Mark S. Klempner, Yang Wang

## Abstract

**Background:** Enterotoxigenic *Escherichia coli* (ETEC) cause diarrheal illness in infants in the developing world and travelers to endemic countries including military personnel. ETEC infection of the host involves colonization of the small intestinal epithelium and toxin secretion leading to watery diarrhea. There is currently no vaccine licensed to prevent ETEC. CFA/I is one of the most common colonization factor antigens (CFAs). The CFA/I adhesin subunit, CfaE, is required for ETEC adhesion to host intestinal cells. Human antibodies against CfaE have potential to block colonization of ETEC and serve as an immunoprophylactic against ETEC-related diarrhea.

**Methods:** Mice transgenic for human immunoglobulin genes were immunized with CfaE to generate a panel of human monoclonal IgG1 antibodies (HuMAbs). The most potent IgG1 identified in the *in vitro* functional assays were selected and isotype switched to secretory IgA (sIgA) and tested in animal colonization assays via oral administration.

**Results:** Over 300 unique anti-CfaE IgG1 HuMabs were identified. The lead IgG1 anti-CfaE HuMAbs completely inhibited hemagglutination and blocked adhesion of ETEC to Caco-2 cells. Epitope mapping studies revealed that HuMAbs recognized epitopes in the N-terminal domain of CfaE near the putative receptor binding site. Oral administration of anti-CfaE antibodies in either IgG or secretory IgA isotypes inhibited intestinal colonization in mice challenged with ETEC. A two to four log decrease of colony forming units was observed as compared to irrelevant isotype controls.

**Conclusions:** We identified fully human monoclonal antibodies against CfaE adhesion domain that can be potentially employed as an immunoprophylaxis to prevent ETEC-related diarrhea.

## Introduction

*Enterotoxigenic Escherichia coli*, ETEC, is one of the main causes of diarrhea in infants in the developing world as well as the major cause of traveler’s diarrhea (1). Transmission of ETEC occurs when contaminated food or water is ingested. ETEC infections are characterized by diarrhea, vomiting, stomach cramps, and in some cases mild fever. Symptoms usually occur 1-3 days after infection and last for a few days (2). When adult travelers develop ETEC diarrhea, a short course of antibiotics can decrease the duration and volume of diarrhea. However, ETEC strains are becoming increasingly resistant to antibiotics (3–5) and there are currently no licensed vaccines for protecting travelers against ETEC diarrhea.

ETEC mediates small intestine adherence through filamentous bacterial surface structures known as colonization factors (CF). Once bound to the small intestine, the bacteria produce heat-labile toxins (LT) and/or heat-stable toxins (ST) that, through a cascade process, cause a net flow of water from the cell into the intestinal lumen, resulting in watery diarrhea (6, 7). ETEC vaccine development efforts have focused on the induction of host immunity against CFs or toxins using cellular or subunit-based approaches. LT have been considered as possible target based on its strong immunogenicity while ST was not initially considered due to poor immunogenicity and potent toxicity. Progress has been made recently in the identification of effective LT-ST toxoids (8, 9). However, anti-ST and anti-LT antibody responses may not be sufficient for effective protection against ETEC diarrhea (8). Instead, the toxin itself may be useful as adjuvant to improve immunogenicity and efficacy when combined with anti-colonization response (10).

Development of an effective immunoprophylactic against ETEC bacterial attachment and colonization has long been considered as an effective approach to prevent ETEC diarrhea (11, 12). The attachment and colonization steps are critical for bacteria to effectively produce toxin and represent a potential strategic target for preventing ETEC infection. The first human-specific ETEC fimbriae to be described was Colonization Factor Antigen I (CFA/I) (13). CFA/I is one of the most prevalent colonization factor antigens expressed by pathogenic ETEC isolates (14, 15). CFA/I is composed of a minor adhesin subunit (CfaE) at the tip of the fimbriae that stabilizes the structure and a long homopolymeric subunit (CfaB) making up the stalk of the structure.

Recent studies have demonstrated that the adhesin subunit itself can provide sufficient immunity to prevent ETEC adhesion and subsequent infection (16, 17). In animal models, maternal vaccination with CfaE resulted in passive protection of neonatal mice from lethal challenge with the ETEC strain H10407. In human clinical trials, a hyperimmune bovine IgG (bIgG) was generated by immunization of a cow with a recombinant form of CfaE and evaluated as a prophylactive treatment in healthy volunteers challenged with ETEC. Oral administration of bIgG antibodies raised against CFA/I minor pilin subunit, CfaE, led to the protection of over 60% of the test group, suggesting that adhesin-based protective antibodies could be used as immunoprophylaxis against ETEC (16).

Here we describe the identification of a panel of anti-CfaE human monoclonal antibodies (HuMabs) that are active against wild-type (or fully virulent) ETEC with high potency in functional assays. Oral administration of the lead HuMabs in either IgG or secretory IgA, the immunoglobulin predominantly secreted at mucosal surfaces, form led to 1-2 log_10_ decreases in colony forming units in an animal model with ETEC challenge. These anti-CfaE HuMabs have potential to be explored as an oral immunoprophylaxis against ETEC infection.

## Results

### Generation of anti-CfaE HuMabs

The N-terminal portion of the adhesin CfaE acts as the receptor binding domain of CFA/I adhesion to host cells (18). To generate a panel of HuMabs that can provide anti-adhesive immunity, eight mice transgenic for human immunoglobulin heavy and light chain genes (Bristol-Myers Squibb; HuMab mice) were immunized with the N-terminal adhesin domain of CfaE fused to maltose binding protein (MBP-CfaE-N). Serum response to MBP-CfaE-N was measured by ELISA. Spleens from mice with positive ELISA response were harvested and fused to melanoma cells to generate hybridomas. A total of 1895 hybridomas were found reactive to MBP-CfaE-N but not the MBP tag itself. RT-PCR was performed on 900 hybridomas to determine the antibody heavy chain gene sequences. A total of 360 HuMabs with unique sequences were selected for further characterization.

### Selection of ten lead HuMabs in mannose resistant hemagglutination assays

All 360 unique HuMabs were purified and tested for their ability to inhibit mannose resistant hemagglutination of human group A erythrocytes (MRHA). MRHA has long been considered as a surrogate method for assessment of ETEC adhesion to the intestinal mucosa (19). The results of the MHRA assays were reported as the maximal inhibitory concentration (IC_100_). 36 of all 360 HuMabs showed IC_100_ activity in nanomolar concentration range. Ten HuMabs were selected as lead candidates with the IC_100_ values between 0.13-ug/mL and 0.24 ug/mL. The heavy chain and light chain gene regions of lead HuMabs were amplified from hybridoma cells and cloned into an immunoglobulin G1 expression vector for antibody expression and purification as previously described. Heavy and Light variable gene families of the lead HuMabs are reported in Table 1.

**Table 1.** Anti-CfaE Hu-MAb Heavy and Light variable gene families

|  | Heavy chain sequence |  |  | Light chain sequence |  |
| --- | --- | --- | --- | --- | --- |
| clone # | Vh | D | Jh | VI | Jk |
| 68-51 | 4-34 | 2-08 | 3 | 1-12 | 2 |
| 68-61 | 1-69 | 2-21 | 3 | 1-16 | 2 |
| 68-97 | 4-34 | 7-27 | 3 | 1-12 | 2 |
| 68-90 | 4-34 | 7-27 | 3 | 1-12 | 2 |
| 68-75 | 4-34 | 2-02 | 6 | 1-13 | 1 |
| 67-102 | 4-34 | 2-15 | 3 | 1-12 | 4 |
| 840-53 | 30-30 | 7-27 | 4b | 1-06 | 1 |
| 68-48 | 4-34 | 7-27 | 6 | 1-13 | 1 |
| 837-6 | 3-23 | 6-06 | 2 | 1-27 | 5 |
| 68-06 | 1-69 | 4-17 | 3 | 1-16 | 2 |

### Anti-CfaE HuMabs bind to recombinant CfaE and live ETEC strain

ELISA results showed that the concentration-dependent binding to CfaE by the lead HuMAbs was indistinguishable (Figure 1, Panel A). To further differentiate the CfaE-binding activities of HuMabs, antibody affinity was analyzed by surface plasmon resonance using recombinant MBP-CfaE-N. All ten HuMabs showed high affinities to MBP-CfaE-N with dissociation constant (KD) values in the low nanomolar range (0.6 nM to 1.2 nM) (Figure 1, Panel B). HuMab 837-6 showed the highest affinity of the ten with a KD value of 2.3×10^−10^. HuMab 68-51, 68-97, 67-102, 68-48 and 837-6 were found to have higher affinity as compared to HuMab 68-61, 68-90, 840-53 and 68-75.

**Figure 1.**
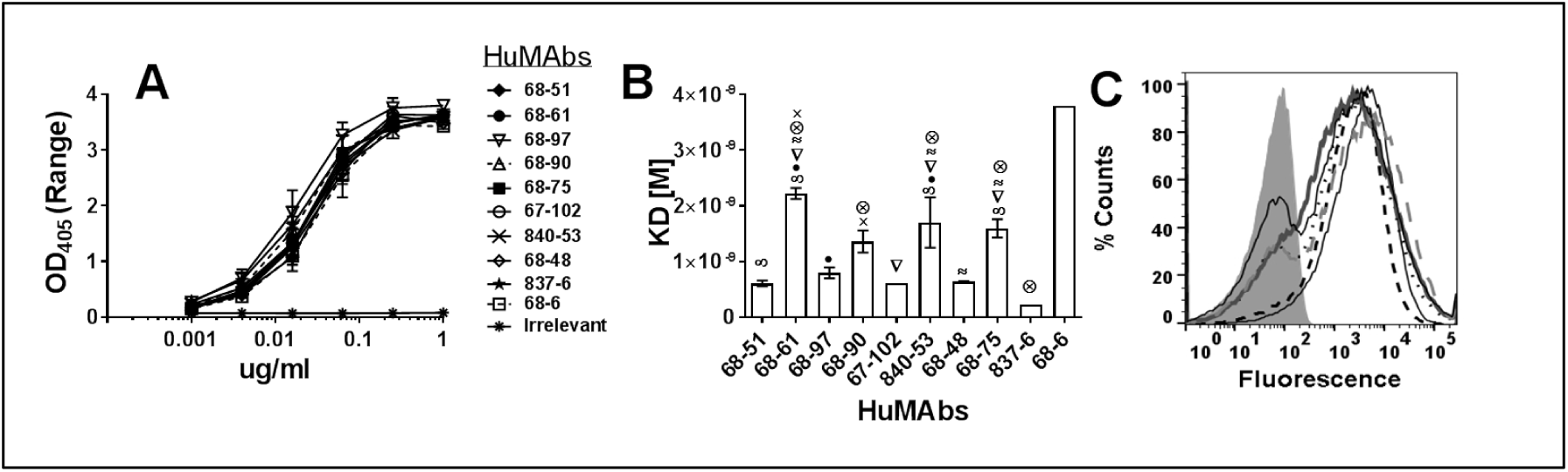
Binding of anti-CfaE MAbs. (A) ELISA. IgG bound to immobilized recombinant CfaE-N was detected with an anti-human IgG Fc-chain-specific alkaline phosphatase-conjugated antibody. Error bars represent the range in OD values observed in two independent experiments. The binding curves of the ten anti-CfaE MAbs are superimposed. (B) Surface plasmon resonance was used to measure the equilibrium dissociation constant (KD). Error bars represent the standard deviation in two independent experiments. All the anti-CfaE antibodies were significantly different compared to the 68–6 HuMAb (P<0.0001). Symbols represent significant differences (P<0.01) between the anti-CfaE HuMAbs using one way ANOVA. (C) Direct binding to live bacterial cells measured by flow cytometry. Gray filled area represents bacteria incubated with an irrelevant antibody.

To assess HuMab recognition of CfaE expressed by live bacteria, H10407 strain was grown in an iron starvation condition to induce CfaE protein expression (20). The bacteria was then incubated with each of the lead ten HuMabs, followed by fluorescence-conjugated secondary antibody and FACS analysis. All HuMabs showed strong binding activity to the H10407 strains. The binding activities were comparable among all ten antibodies (Figure 1, Panel C).

### Anti-CfaE HuMabs prevent ETEC adherence to intestinal cells at low concentrations

To determine whether the lead HuMabs were capable of inhibiting bacterial adhesion, a cell adhesion assay with Caco-2 cells (a human intestinal epithelial cell line) was performed. An example of concentration-dependent inhibition curve is reported in Figure 2, Panel B. The minimal inhibitory concentrations needed to prevent 50% (IC_50_) of bacterial adhesion were reported as antibody potency. All ten HuMabs showed strong potency to block bacteria adhesion at IC_50_ concentrations between 0.3 to 1.3 ug/mL. HuMab 68-51, 68-61 and 68-97 were found to have the lowest IC_50_ values (Figure 2, Panel C). Interestingly, HuMabs showing comparable activities in MRHA assays (Figure 2, Panel A) were more variable in their activities in Caco-2 cell adhesion assays.

**Figure 2.**
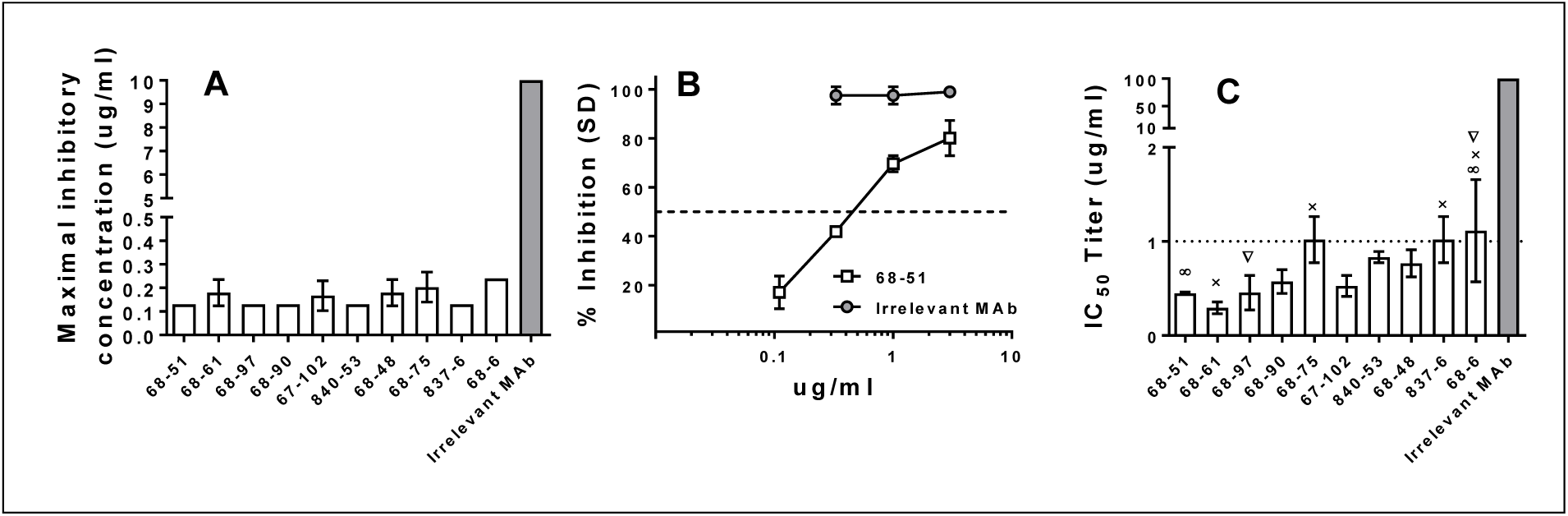
*In vitro* functional activity of anti-CfaE MAbs. (A) Hemagglutination assay. The ability of the MAbs to prevent hemagglutiantion is reported as maximal inhibitory concentration (IC_100_). Error bars represent the standard deviation observed in three independent experiments using different blood donors. (B) Caco-2 adhesion assay. Example of inhibition curve obrtained with MAb 101 and an irrelevant control. (C) The minimal effective IgG dose to prevent 50% (IC_50_) of bacterial adhesion to intestinal Caco-2 cells was used to determine antibody potency ranking. Error bars represent the standard deviation in three to four independent experiments. All the anti-CfaE HuMAs were significantly different compared to the irrelevant MAb (P<0.0001). Symbols represent significant differences (P<0.01) withing the anti-CfaE HuMAbs using one way ANOVA.

### Epitope Mapping of lead anti-CfaE HuMabs

To define the antibody-binding epitope, putative antibody-antigen interaction models were established based on a previously resolved CfaE structure (PDB ID 2Hb0) and the lead HuMab antibody sequences using an antibody modeling program, BioLuminate (Schrodinger). This software suite develops models of antibody structures from their sequences, followed by computational docking to identify high-confidence antibody-antigen complex models. Based on these models, the software identified potential residues critical for binding interaction. The effect of these residues on the binding activity of the HuMAbs was analyzed by experimental alanine scanning followed by ELISA. ELISA results indicated that mutating five of the predicted residues to alanine affected HuMAb binding (Figure 3). The R67A mutation eliminated binding activity of HuMab 68-51 and 68-97 (Panel B), while the Y183A mutation affected binding activity of HuMab 68-51, 68-97, 68-90, 67-102 and 840-53 (Panel E). R145A mutation abolished binding activity of HuMab 837-5 (Panel D). T91A mutation eliminated binding activity of HuMAb 840-53 and reduced binding activities of HuMAbs 68-51, 68-61, 840-53 and 68-48 (Panel C). N127A mutation eliminated binding activity of HuMAb 68-61 and reduced binding of HuMAbs 68-48 and 68-6 (Panel F). Summary of the residues discovered to affect binding are shown in Figure 3, Panel G. All mutations were found on the surface exposed loops of the N-terminal domain of the CfaE (Figure 3, Panel H). No residues involved in the binding of MAb 68-75 to CfaE were identified.

**Figure 3.**
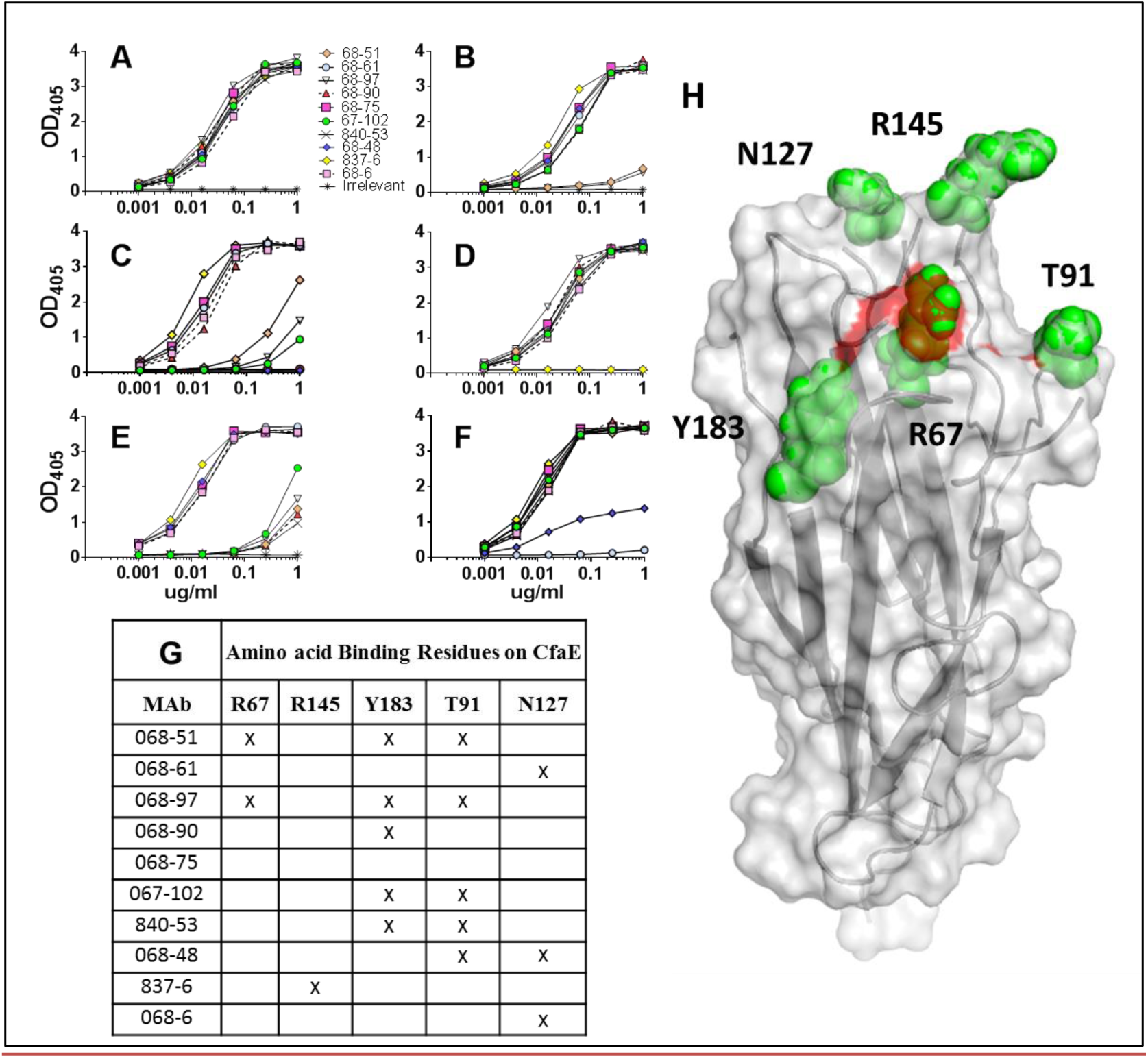
Epitope mapping studies. Binding of anti-CfaE MAbs to mutants of recombinant CfaE as measured by ELISA. (A) Wild-type CfaE. (B) Arg67Ala mutant. (C) Thr91Ala mutant. (D) Arg145Ala mutant. (E) Tyr183Ala mutant. (F) Asn127Ala mutant. (G) Crystal structure of N-terminal CfaE molecule with the five residues involved in the anti-CfaE MAbs binding showed as green spheres. Highlighted in red the three arginine forming the putative receptor binding domain.

#### Isotype switch of anti-CfaE HuMabs to sIgA

Seven IgG1 HuMab (68-51, 68-61, 68-97, 68-90, 67-102, 68-48, and 840-53) found to have the lowest IC_50_ values in Caco-2 cell adhesion assays were selected as the leads for further characterization in immunoglobulin class switching. Antibody variable regions were cloned into an expression vector with IgA constant region to generate monomeric IgA. Monomeric IgA antibodies were also co-expressed with J chain with or without secretory component to produce dimeric IgA (dIgA) and secretory IgA (sIgA) respectively. Antibodies with various isotypes were tested for their functionality in Caco-2 cell adhesion assays (Figure 4). In general, all the antibodies retained or increased *in vitro* functional activity when converted into dIgA or sIgA. Specifically, *in vitro* functional activity of 68-61 was not altered significantly when converted to either dimeric or secretory IgA molecule. In contrast, Ig class switching to either dimeric or secretory IgA forms caused significantly improvement of functional activity for HuMAbs 68-97, 840-53 and 68-48. Interestingly, HuMAb 68-90 only saw a significant improvement when Ig class switch to dimeric or secretory IgA1. Additionally, conversion from an IgG1 to a dimeric IgA1 or IgA2 did not affect functional activity of HuMAb 67-102, but switching from dimeric to secretory IgA1 or IgA2 did significantly increase *in vitro* activity of 67-102. Due to low expression yields, we were not able to generate sufficient 68-51 sIgA1 or sIgA2 for *in vitro* testing.

**Figure 4.**
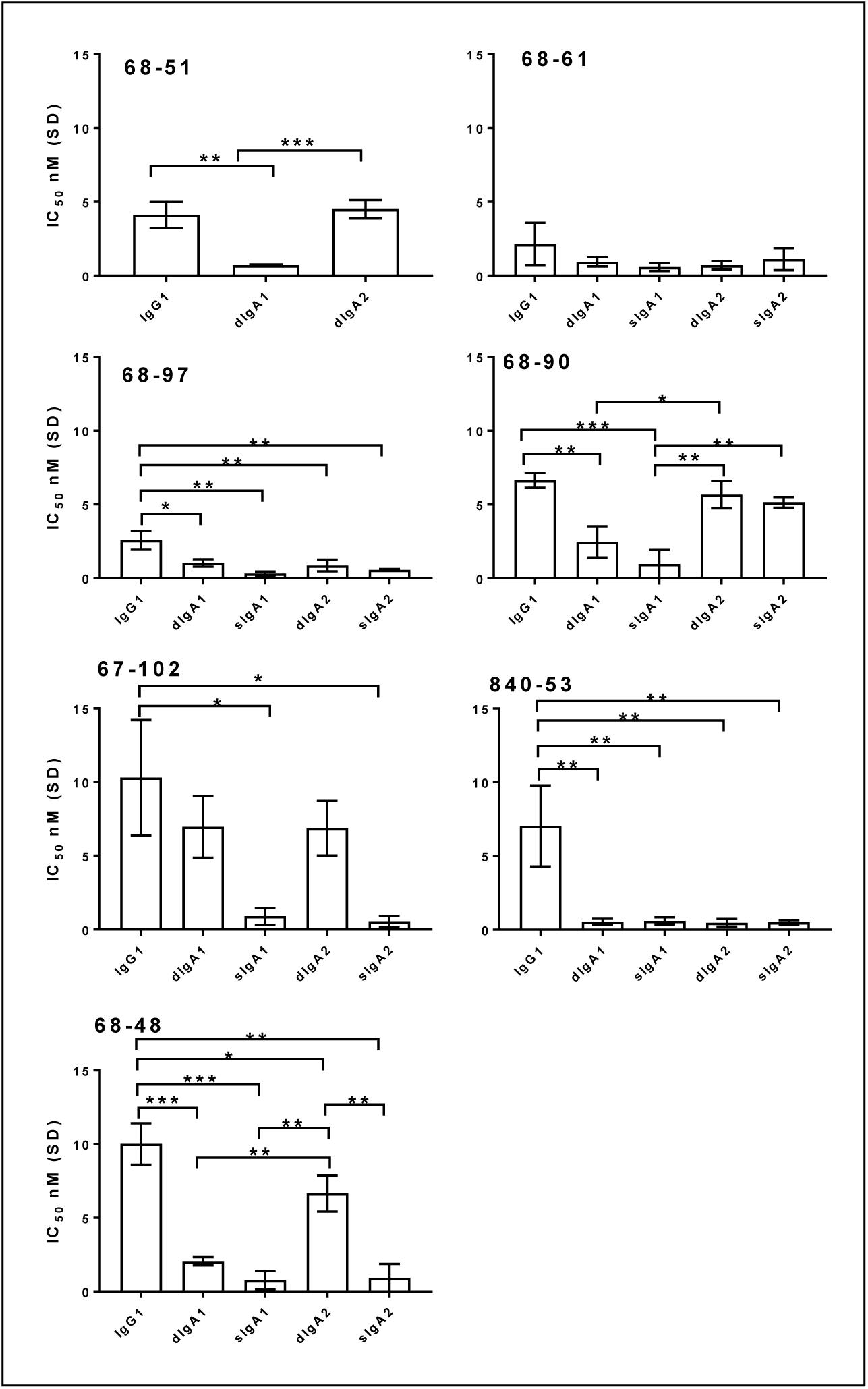
Ig Class switching of anti-CfaE MAbs. Caco-2 adhesion assay. The minimal effective IgG dose to prevent 50% (IC_50_) of bacterial adhesion to intestinal Caco-2 cells was used to determine antibody potency ranking. Error bars represent the standard deviation in three to four independent experiments. *P < 0.01, **P < 0.001, ***P < 0.0001.

### Anti-CfaE HuMabs prevent ETEC colonization in the small intestine of a mouse model

HuMAbs 68-61, 68-97, and 840-53 were found to have the lowest IC_50_ values as IgG1, sIgA1, and sIgA2 and were selected as the leads for further characterization in animal studies (Figure 5). Groups of 5 DBA2 mice were given a mixture of bacteria and anti-CfaE HuMAbs (10 mg/Kg) by oral gavage. 24 hours after challenge, the mice were euthanized and the CFU in the small intestine were counted as described in the methods. The efficacy of the anti-CfaE HuMAbs was assessed by determining whether the HuMAbs could prevent adhesion of bacteria to the small intestine compared to an irrelevant isotype control. In the 68-61 group, treatment with IgG1 decreased CFU by100 fold compared to the irrelevant antibody control. A similar result was observed for 68-61 sIgA2 and sIgA1 compared to the irrelevant control. The reduction of CFU observed in the 68-97 group compared to the irrelevant control was similar across the different subclasses. In the 840-53 group, mice treated with IgG1 showed less bacteria compared to sIgA2, while sIgA1 also showed a decrease in bacteria relative to sIgA2, though these differences were not significant.

**Figure 5.**
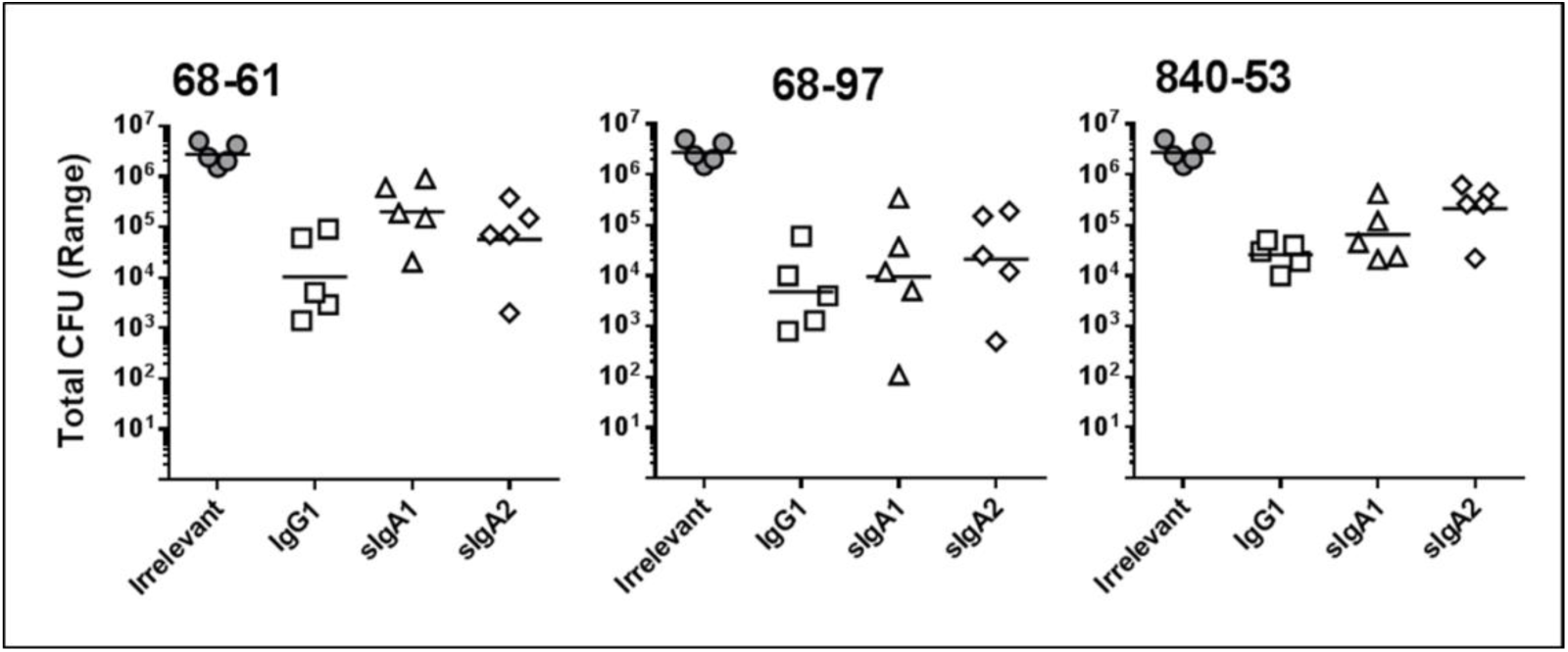
In vivo studies. DBA/2 mice were challenged intra-gastrically with 10^7^ CFU pre-incubated with 10 mg/Kg of HuMAbs. Animals were euthanized 24 hours after challenge and bacterial colonies in the small intestine were counted. Five animals were tested for each condition. All the anti-CfaE HuMAbs were significantly different compared to the irrelevant MAb (P<0.001).

## Discussion

Recent studies from our laboratory and other demonstrated the feasibility of using HuMAbs for pre-exposure prophylaxis against infectious diseases (21). A similar strategy could be used to prevent diarrhea disease caused by enteric pathogens such as ETEC. ETEC fimbriae adhesin subunits mediate adherence of the bacteria to the small intestine as the fist step of a cascade of events leading to diarrhea. Studies in animal models and in human vaccine clinical trials have demonstrated that immunity against adhesion is sufficient to elicit effective protection against ETEC infection (16, 17). In the present study, we produced and characterized HuMabs that recognize N-terminal portion of CfaE (CfaE-N), the adhesin subunit involved in bacterial adhesion to host cells. Among the three hundred isolated HuMabs, ten lead candidates were selected based on their ability to block mannose-resistant hemagglutination (MRHA) and inhibit adhesion of ETEC to Caco-2 cells. When administrated orally to mice by pre-mixing with ETEC, the selected lead HuMabs significantly decrease colony formation in the small intestine.

Structural analysis have revealed a positively charged binding pocket in the N-terminal domain of CfaE responsible for ETEC interaction with the negatively charged sialic acid of the host epithelial cells (shown as red surface in Figure 3, Panel H). The binding pocket is formed by three arginine residues, Arg67, Arg181 and Arg182, and a cluster of surrounding residues, which are all highly conserved among the Class-5 fimbriae adhesins. Mutations of the arginine residues result in a complete loss of binding activity of ETEC to host cells (18). Putative antibody-antigen interaction models were established utilizing the sequences of CfaE (PDB ID 2HB0) and the anti-CfaE HuMAbs with the aid of an antibody modeling program (BioLuminate, Schrödinger). A set of residues predicted to be important for the binding interaction based on the top-ranked model were selected for experimental alanine scanning, and were confirmed to be energetically critical using ELISA. HuMabs 68-51 and 68-97, the antibodies with the highest *in vitro* functional activity, were found to bind directly to Arg67 (part of the putative receptor binding domain). Four more residues, Arg145, Thr91, Tyr183, and Asn127, located in the proximity of the receptor binding domain, were recognized by other nine lead anti-CfaE HuMabs (Figure 3). Together, our data suggest that the protective HuMabs are likely to be elicited by epitopes within or near the positively charged binding pocket. HuMabs binding to ETEC either directly to the putative receptor binding domain or its proximity region may result in the blockage of ETEC and host cell interactions. Unfortunately, we didn’t identify any residues involved in the binding of HuMAb 68-75 to CfaE, for which a substitution different from Alanine maybe required in order to elucidate the location of the epitope recognized by this MAb.

The development of novel immunoprophylaxis to prevent disease depends on the availability of animal models that faithfully recapitulate the disease in human. This is particularly the case for development of immnoprophylaxis against ETEC since the colonization factors are highly species specific. In a recent study, a neonatal mouse model using DBA/2 mice was reported to be effective in investigating a adhesin based vaccine. A lethal dose was determined and the pups were rescued by maternal vaccination or by oral administration of hyperimmune anti-CFA/I and anti-CfaE bovine colostral IgG (16). To evulate the potency of lead HuMabs, we have tried to established the same model described by Luiz et al. However, maternal aggression and high rates of pup cannibalism limited our cability to succesfuly conduct the assay. Alternatively, we established an adult colonization mouse model using DBA/2 mice. Six week old mice were challenged with a non-lethal dose of ETEC that would lead to a consistant level of bacteria burden in the small intestine as described on other studies (22, 23). The ability of HuMab to inhibit adherence of ETEC was measured as a reduction of colony count compared to an irrelevant iotype control.

IgG and sIgA are both present in the small intestine as effector molecules of mucosal immune system. sIgA is considered to be one of the most important effector molecules because it constitutes the primary immune defence against pathogens at the mucosal surface (24). In secretory IgA, two IgA monomers are covalently linked by a joining chain (J-chain), and stabilized by a polypeptide called the secretory component that make the molecule more resistant to digestion in the small intestine than IgG (25). Early studies also have suggested that the secretory component may have its own antimicrobial activity to block epithelial adhesion of enterotoxigenic E. coli (26). To assess the funtional differences between IgG and sIgA, three lead HuMabs, 68-61, 840-53 and 68-97 were selected and class switched to IgA1 and IgA2 in both dimeric and secretory forms. The antibody activities *in vitro* were generally well preserved after the class switch. Pre-incubation of bacteria with either sIgA or IgG forms lead to significant reduction of CFU in the small intestine compared to the irrelevant controls (P<0.001). 68-61 and 840-53 IgG1 showed greater activity than their sIgA1 and sIgA2 forms respectively, but the differences were not significant. No significant diferences were observed in the 68-97 group. Overall, class switch to sIgA did not show benefits over the IgG form, possibly due to the limitation of our method to assess human antibody stability in the mouse intestine.

In conclusion, we identified a panel of HuMabs that potently inhibit ETEC binding to intestinal epithelia cells *in vitro* and *in vivo*. Additional pre-clinical studies are planned to desmonstrated the safety and efficacy in a *Aotus* monkey model with ETEC challenge (27, 28). To date, over 25 human-specific ETEC Colonization Factor Antigens (CFAs) types have been described and most CFAs are fimbrial proteins (29). Eight common fimbriae (CFA/I, CS4, CS14, CS1, CS17, CS19, PCF071 and CS2) belonging to Class-5 are expressed by pathogenic strains that cause the majority of moderate to severe ETEC diarrheal cases (13, 14, 30). Anti-CfaE monoclonal antibodies have been previously shown to cross-react and cross-protect with heterologus CFAs of the Class-5 fimbriae (31). For this reason, further investigation is needed to explore the cross-protectivity of the anti-CfaE HuMab against heterologus CFA ETEC strains. In the absence of a vaccine for ETEC, our study provide the first proof of concept that oral administartion of protective antibody could potentially be an effective strategy for prophylaxis against ETEC.

## Methods

### ETEC test strains

H10407 expressing CFA/I fimbriae was purchased from ATCC (ATCC^®^ 35401™). ETEC strain H10407 was cultured on 2% agar containing 1% Casamino Acids (Sigma) and 0.15% yeast extract (Fisher Bioreagents) plus 0.005% MgSO_4_ (Sigma) and 0.0005% MnCl_2_ (Sigma) (CFA agar plates) overnight at 37°C. 1×10^8^ colony forming units/mL were resuspended in 20% glycerol (Sigma) in PBS solution, and kept frozen at −80°C until needed.

#### Antigen cloning, expression, purification

The nucleic acid sequences of N-terminal adhesin domains of CfaE (GenBank M55661) was cloned into a pMAL-C5X vector (Addgene) in-frame with a MBP tag to express as periplasmic proteins with improved solubility (MBP-CfaE-N).

The donor strand complement was included to ensure the overall protein expression and stability as reported (32). All cloned constructs were transformed into SHuffle^®^T7 Competent *Escherichia coli* (NEB), and expression was induced with 1mM IPTG. Bacteria were lysed, and proteins were purified with amylose resin (NEB) and eluted with 20 and 50 mM Maltose (Sigma).

#### Mouse Immunization, Hybridoma Generation, and Antibody Cloning

Transgenic mice containing human immunoglobulin genes and inactivated mouse heavy and κ light chain genes (Bristol-Myers Squib) were immunized with 50 μg of MBP-CfaE-N weekly with the Sigma adjuvant system (Sigma) for 6-10 weeks. Anti-CfaE titer in mouse serum was measured by enzyme-linked immunosorbent assay (ELISA). Hybridomas were generated following a standard PEG fusion protocol (21). Hybridoma supernatants were screened for reactivity to MBP-CfaE-N, and positive cell clones were selected for antibody sequencing. The heavy chain and light chain variable regions were amplified from hybridoma cells and cloned into two pcDNA 3.1 (Thermo Fisher) vectors containing k light constant and IgG1 heavy constant chain respectively as previously described (21).

### IgA class switching

Primers were designed to amplify the variable heavy chain of each IgG antibody, and products were digested and ligated into a pcDNA 3.1 vector containing heavy constant IgA1 and IgA2 chains. Each vector was transformed in NEB5-α competent cells and sequences were verified ahead of transient transfection. In order to get dimeric IgA, the heavy and light chain vectors were co-transfected with pcDNA containing DNA for the connecting J-Chain using an ExpiCHO expression system (Life Technology). For secretory expression, a pcDNA containing secretory component was added to the transfection reaction in a 1:1 ratio. Supernatant was run through a column of CaptoL resin to capture the light chains of antibodies (GE Life Sciences). Antibodies were dialyzed against phosphate buffered saline before moving into size exclusion chromatography to separate out the desired dimeric or secretory antibodies using a HiLoad 26/600 Superdex 200 pg size exclusion column (GE Healthcare Life Sciences). Desired fractions were pooled, concentrated and quality tested by SDS-PAGE and western blots (Figure S1).

#### ELISA assay

For binding activity of purified HuMAbs against CfaE, 96-well plates (Nunc) were coated overnight at 4°C with 2 μg/mL of purified MBP-CfaE-N. Plates were blocked with 1% BSA + 0.05% Tween 20 in PBS. Purified HuMabs were diluted in 1× PBS + 0.1% Tween 20 and added to plates for 1 hour. Plates were stained with alkaline phosphatase-conjugated goat anti-human IgG Fcγ (Jackson ImmunoResearch Laboratories) (1:1,000) for 1 hour and developed using p-nitrophenyl phosphate (ThermoFisher Scientific). Absorbance at an OD of 405 nm was measured on an Emax precision plate reader (Molecular Devices).

#### SPR analysis

Surface plasmon resonance (SPR) technology was used to assess the binding properties of the HuMAbs (Biacore T200 instrument; GE Healthcare). A total of 2,700 response units (RU) of anti-human IgG MAb (human antibody capture kit; GE Healthcare) was coupled to a CM5 sensor chip using standard amine coupling chemistry. In multi-cycle kinetics experiments, 25 to100 RU of each anti-CfaE HuMAb was captured on the anti-human IgG Mab bound sensor chip. Various concentrations of soluble recombinant MBP-CfaE-N antigen ranging from 1.56 nM to 50 nM were injected over the chip surface at a flow rate of 30 μl/min. An association step of 60s was followed by a dissociation step of 180s, and the final dissociation step was 600s. Regeneration of the sensor chip surface was accomplished using 3M MgCl_2_. Experiments were performed at 25°C. Kinetic data were analyzed using Biacore T200 Evaluation (version 3.0) software and a 1:1 binding model. All chemicals for the Biacore experiment were purchased from GE Healthcare.

#### Flow cytometry

Binding of the HuMAbs to the surface of live bacteria was measured by flow cytometry as described previously (33). H10407, which expresses the target CFA/I antigen, was used as the test strain. Briefly, bacteria were grown in CFA medium supplemented with 50 μM deferoxamine overnight at 37°C with gentle shaking (20). To measure HuMAb binding, a fixed concentration of anti-CfaE HuMAb (10 μg/mL) or, as a negative control, 100 μg/mL of an irrelevant MAb, was incubated with 10^7^ bacteria/mL. Bound antibody was detected using CF488-conjugated goat anti-human IgG (Biotium).

### Mannose resistant hemagglutination assay of human group A erythrocytes

ETEC cultures were taken from frozen cell banks and diluted in sterile 0.15 M saline solution until reaching an OD_600nm_ of 1 for the assay. Human erythrocytes type A+ stored in K3EDTA were washed three times with 0.15 M saline solution and resuspended in the same solution to a final concentration of 1.5% (vol/vol). In a U-bottom 96-well plate (Nunc Thermo Scientific) 100 μl of HuMAb was added in duplicate to the top row and diluted 1:2 down the plate in 0.15 M saline solution. 50 μl of appropriately diluted ETEC was added to each well together with 50 μl of 0.1 M D-mannose solution (sigma). The plate was incubated for 10 minutes at room temperature. Afer incubation, 50 μl of blood solution was added to the plate and mixed well (200 μl final volume). Plates were allowed to sit stagnant at 4°C for two hours. Hemagglutination was then observed without the aid of magnification. The absence of a pellet of red blood cells at the bottom of the well is indicative of positive hemagglutination. Blood was ordered fresh every week (BioreclamationIVT).

#### Caco-2 adhesion assay

Caco-2 cells seeded at 1 × 10^5^ cells/mL were grown in 24-well tissue culture plates containing Dulbecco's modified Eagle's medium (DMEM), at 37°C in 5% CO_2_ static. Frozen bacterial banks were streaked on CFA agar plates and grown overnight at 37°C. The next day, bacteria were resuspended in PBS and diluted until reaching an OD_600nm_ of 0.1. HuMab dilutions were set up in a deep well plate. Antibody dilutions and bacteria were combined in a 1:10 ratio and allowed to shake at 300 rpm for one hour at room temperature. After incubation, 0.2 mL of antibody/bacteria mixture was added to each well containing Caco-2 cells. The cells were then incubated statically for 3 hours at 37°C. Cells were then washed four times with 1 mL PBS to remove non-adherent ETEC cells.

Afterwards, Caco-2 cells were dislodged with 0.2 mL 0.25% trypsin. Cells were collected via gentle centrifugation and resuspended in 1mL PBS. Dilutions were plated on CFA agar plates and colonies counted the next day. IC_50_ was defined as concentration of HuMAb needed to inhibit 50% of ETEC adhesion to the Caco-2 cells, compared to an irrelevant isotype antibody.

#### Animal assays

6-8 week old DBA-2 mice were pretreated with streptomycin (5 g/L) in the drinking water for 24-48 hours. Twelve hours prior to bacteria administration the water was replaced with regular drinking water. One hour prior to bacteria administration, mice received cimetidine (50 mg/kg) i.p. to reduce the effect of stomach acid on ETEC. A total of 10^7^ CFU of H10407 ETEC strain diluted in PBS were incubated with 10 mg/Kg of an anti-CfaE HuMAb or an irrelevant MAb (Purified human secretory IgA, MP Biomedicals) one hour prior challenge. Bacteria and HuMAbs were administered in 200 μl volume by oral gavage using 20 g bulb-tip feeding needles. The mice were allowed to survive for 24 hours. 12 hours prior euthanasia food was withdrawn. Following isolation of the small intestine, two segments of ileum (3 cm each), beginning within 0.5 cm of the ileocecal junction and extending proximally 6 cm, were removed and placed in 1mL sterile PBS (22). Tissues were mechanically homogenized. Samples were serially diluted on MacConkey agar plates and incubated overnight at 37°C. Bacterial CFUs were counted the next day. To confirm that recovered bacteria were the inoculum strain, bacterial colonies grown on culture plates were routinely tested by PCR using specific primers (22), which flank the eltAB operon encoding the LT holotoxin of H10407.

#### Epitope mapping

Bioluminate software (Schrödinger) was used to identify CfaE residues involved in antibody-antigen recognition. A total of 22 amino acids predicted by the software to be involved in the interaction between anti-CfaE HuMAbs and the N-terminal portion of CfaE were individually mutated to Alanine using BioXp™ 3200 System (SGI-DNA). The genes were cloned into pMAL-C5x vector and the resulting 22 constructs were transformed, expressed and purified as described above. An ELISA assay was performed to determine binding of the HuMAbs to the mutant proteins compared to the wild-type.

### Statistical analysis

Statistical calculations were performed using the software Prism version 7.03 (GraphPad Software, La Jolla, CA). Comparisons between the hemagglutination or Caco-2 titers of respective antibodies were performed using multiple comparisons, Bonferroni test, one way ANOVA.

